# Expression of Sonic Hedgehog and Pathway Components in the Embryonic Mouse Head: Anatomical Relationships Between Regulators of Positive and Negative Feedback

**DOI:** 10.1101/2021.03.30.437697

**Authors:** Crystal L. Sigulinsky, Xiaodong Li, Edward M. Levine

## Abstract

**Objective:** The Hedgehog pathway is a fundamental signaling pathway in organogenesis. The expression patterns of the ligand *Sonic Hedgehog* (*Shh*) and key pathway components have been studied in many tissues but direct spatial comparisons across tissues with different cell compositions and structural organization are not common and could reveal tissue-specific differences in pathway dynamics.

**Results:** We directly compared the expression characteristics of *Shh*, and four genes with functional roles in signaling and whose expression levels serve as readouts of pathway activity in multiple tissues of the embryonic mouse head at embryonic day 15.5 by serial *in situ* hybridization. The four readout genes were the positive feedback regulator *Gli1*, and three negative feedback regulators, *Patched1, Patched2*, and *Hedgehog Interacting Protein*. While the relative abundance of *Gli1* was similar across tissues, the relative expression levels and spatial distribution of *Shh* and the negative feedback regulators differed, suggesting that feedback regulation of hedgehog signaling is context dependent. This comparative analysis offers insight into how consistent pathway activity could be achieved in tissues with different morphologies and characteristics of ligand expression.

## INTRODUCTION

Shh is a secreted glycoprotein belonging to the Hedgehog (Hh) family of intercellular signaling molecules. The mechanics and regulation of Hh signaling are complex extending from ligand production through signal transduction to the cell- and tissue-specific responses (reviewed in (1-4)). In its simplest iteration (Fig. 1A), binding of Shh to the PATCHED receptors, Patched 1 (Ptch1) or, in some cases, Ptch2, relieves inhibition of the G-protein coupled receptor Smoothened (Smo). Activated Smo inhibits proteolytic processing of the GLI transcriptional effectors Gli2 or Gli3 into truncated repressor forms through destabilization of complexes between Gli2 or Gli3 with Suppressor of Fused (Sufu). The resulting accumulation of full-length GLI proteins in the nucleus promotes both the derepression and activation of Hh target genes.

**Figure 1:**
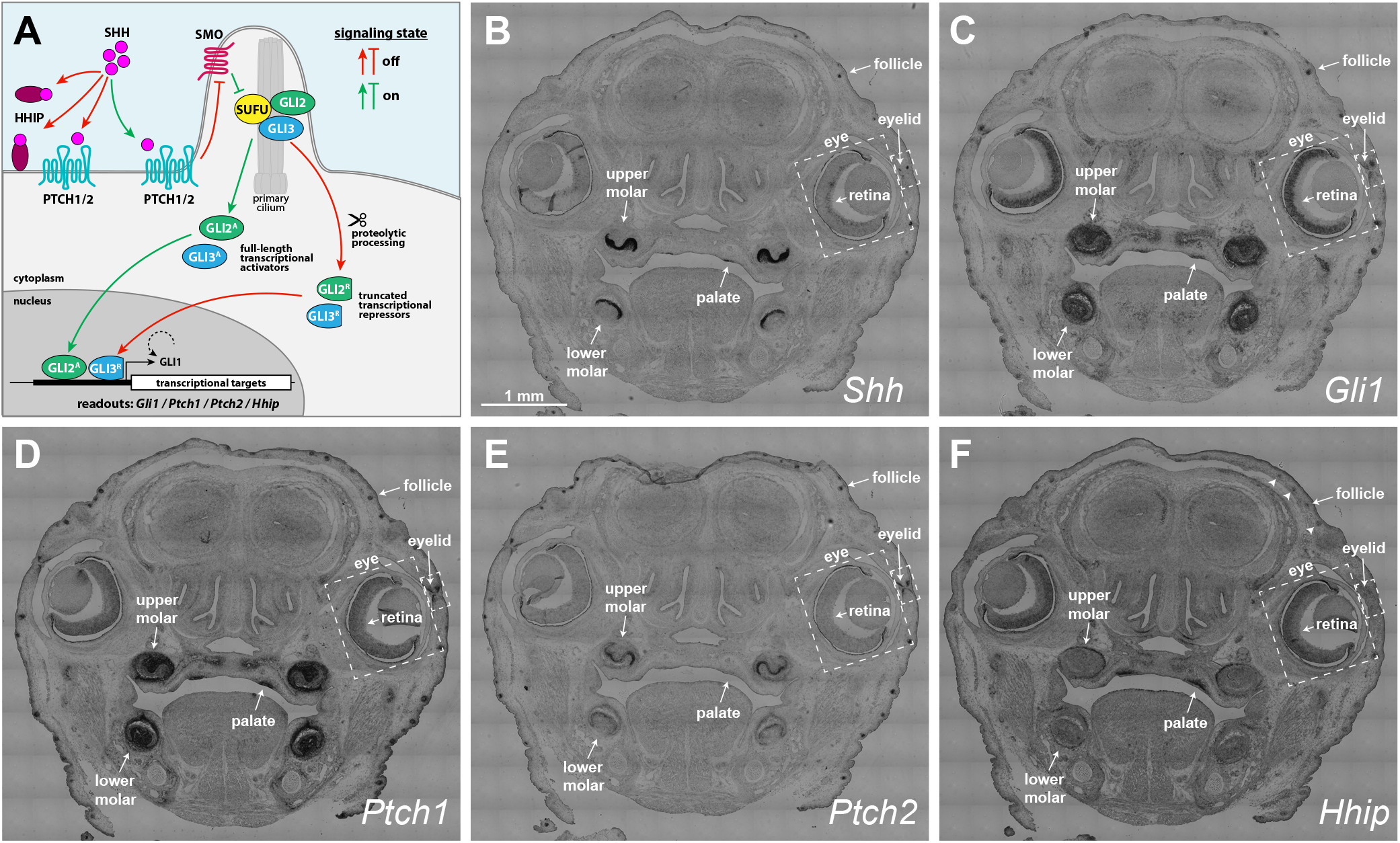
Expression patterns for Shh and Hh pathway components in developing organs of the embryonic mouse head. (A) Simplified schematic of Hh signaling. See Introduction for details. (B-F) *in situ* hybridizations for *Shh* (B), *Gli1* (C), *Ptch1* (D), *Ptch2* (E), and *Hhip* (F) in adjacent coronal sections of the mouse at E15.5. Arrowheads in F denote additional hair follicles.

Transcriptional targets of the Hh pathway not only mediate cellular responses to Hh ligands, but also participate in feedback loops that further regulate Hh pathway activity. The principal positive feedback loop involves the transcriptional effector Gli1. *Gli1* expression is activated in response to Gli2 mediated transduction of Hh signals (5-7).This, together with its activator function, allows Gli1 to increase signaling levels within responding cells while retaining dependence on active Hh signaling. *Gli1* expression is therefore an excellent indicator of Hh pathway activity.

Ptch1, Ptch2, and Hhip participate in negative feedback that act at the level of Hh reception (8-10). *Ptch1* transcripts are also upregulated in response to Hh signaling (11-13), and evaluation of phenotypes and Hh pathway activity in *Ptch1* mutant mice shows that Hh activity is sensitive to *Ptch1* gene dosage (14-16). In addition to Smo inhibition, upregulation of *Ptch1* (*Ptc* in *Drosophila*) also sequesters Hh ligands and desensitizes the cell to Hh signal (17). *Patch2* shares sequence homology with *Ptch1*, binds Hh ligands with high affinity and inhibits Shh-induced changes in gene expression (18, 19). *Ptch2* is also upregulated in response to Hh signaling, but this can be context dependent (18-20). Additionally, Ptch2 fails to block changes in gene expression induced by a constitutively active form of Smo and is unable to replace Ptch1 function in *Ptch1* mutant basal carcinoma cells but does preserve some ligand dependent signaling in *Ptch1-*null fibroblasts (19, 21, 22). Interestingly, Ptch1 and 2 were recently shown to non-autonomously inhibit Smo, possibly through secretion of a cholesterol precursor (10). Like *Ptch1* and *Ptch2, Hhip* is upregulated in response to Hh signaling. Hhip also binds Hh ligands with high affinity and can attenuate Hh signaling through ligand sequestration (8, 9, 23, 24). Thus, like Ptch1 and Ptch2, Hhip also negatively regulates the level of Hh ligands to which the responding cell is exposed.

In our studies on Hh signaling in embryonic retinal neurogenesis in mice, which begins at ∼E11.5, we’ve observed that *Shh* expression can be difficult to detect even though Hh signaling has essential functions in retinal development, and *Shh*, expressed in retinal ganglion cells (RGCs), is the sole Hh ligand employed during this time (reviewed in (25)). We asked, if *Shh* expression was lower in the retina than in other anatomical structures, how would *Gli1* and the expression of the negative feedback regulators compare? And if differences in expression exist across structures, could anatomical differences correlate with how *Shh* and the feedback components are expressed? To address these questions, we performed *in situ* hybridizations for *Shh, Gli1, Ptch1, Ptch2*, and *Hhip* on adjacent serial sections of an E15.5 embryonic mouse head. Through coronal sectioning, direct comparisons were made for each gene across 6 tissues and for the 5 genes in each tissue.

## MAIN TEXT

## Materials and methods

### Animals

129/Sv mice were purchased from The Jackson Laboratory (Bar Harbor, ME, USA). Mice were bred overnight and noon on the day of vaginal plug was considered embryonic day 0.5 (E0.5).

### In situ hybridization

Heads were fixed overnight at 4 °C in 4% formaldehyde in PBS pH7.5, 2 mM EGTA. Fixed tissue was cryoprotected with 20% sucrose/PBS, embedded and frozen in OCT. Adjacent 12 µm serial sections were stained with digoxigenin-labeled anti-sense probes produced by *in vitro* transcription of sequence-verified linearized plasmids (Supplemental Fig. 1). Section in situ hybridization was performed as previously described (26-28).

### Image capture and analysis

Sections were imaged at 10X magnification on a Leica DMR microscope under brightfield illumination. Image tiles (8-bit, 1388 x 1036 pixel) were acquired with a QICAM Fast 1394 (QImaging, Burnaby, Canada) and automated scanning stage (Märzhäuser Wetzlar GmbH, Wetzlar Germany). Mosaic images were assembled using a Syncroscan montaging system (Synoptics Inc, Frederick, MD). Close up views of the hair follicles were imaged at 20X magnification on a Nikon E-600 microscope using differential interference contrast. Images were acquired with a Spot-RT camera (Diagnostic Instruments Inc., Sterling Heights, MI). Due to their small size, hair follicles could not be analyzed for all probes on adjacent serial sections. Thus, similar positions within representative morphologically matched follicles were imaged. Figure assembly was done Photoshop and Illustrator CC (Adobe Systems Inc., San Jose, CA, USA).

## Results

Figure 1 shows the expression patterns of *Shh, Gli1 Ptch1, Ptch2*, and *Hhip* in the context of the head. We identified the upper and lower molars, palatal rugae, retina, eyelid, and hair follicles as tissues for comparison based on the presence of both *Shh* and *Gli1. Shh* expression identified the cellular sources of Hh signal and was most readily detected in the molars (Fig. 1B). Despite the small sizes of the hair follicles, palatal rugae, and eyelid, *Shh* expression was still evident at this scale. In contrast, the retina exhibited a low level of *Shh* expression that was disproportionate to its relatively large size. *Gli1* expression, the indicator of Hh signaling, was similarly robust across all 6 tissues (Fig. 1C). *Ptch1* expression was also robust in all 6 tissues (Fig. 1D) although its expression in the retina appeared lower by comparison to the levels of *Gli1* in each tissue. This is more easily observed in Figure 2. The expression patterns of *Ptch2* were most similar to *Shh*, although expression levels in the retina and palatal rugae were too low to assess at this scale (Fig. 1E). *Hhip* was detected in the molars, hair follicle, palatal rugae, and retina; expression in the eyelid was too low to assess. Interestingly, *Hhip* was abundant in the retina and palatal rugae (Fig. 1F), where *Ptch2* was lowest.

**Figure 2:**
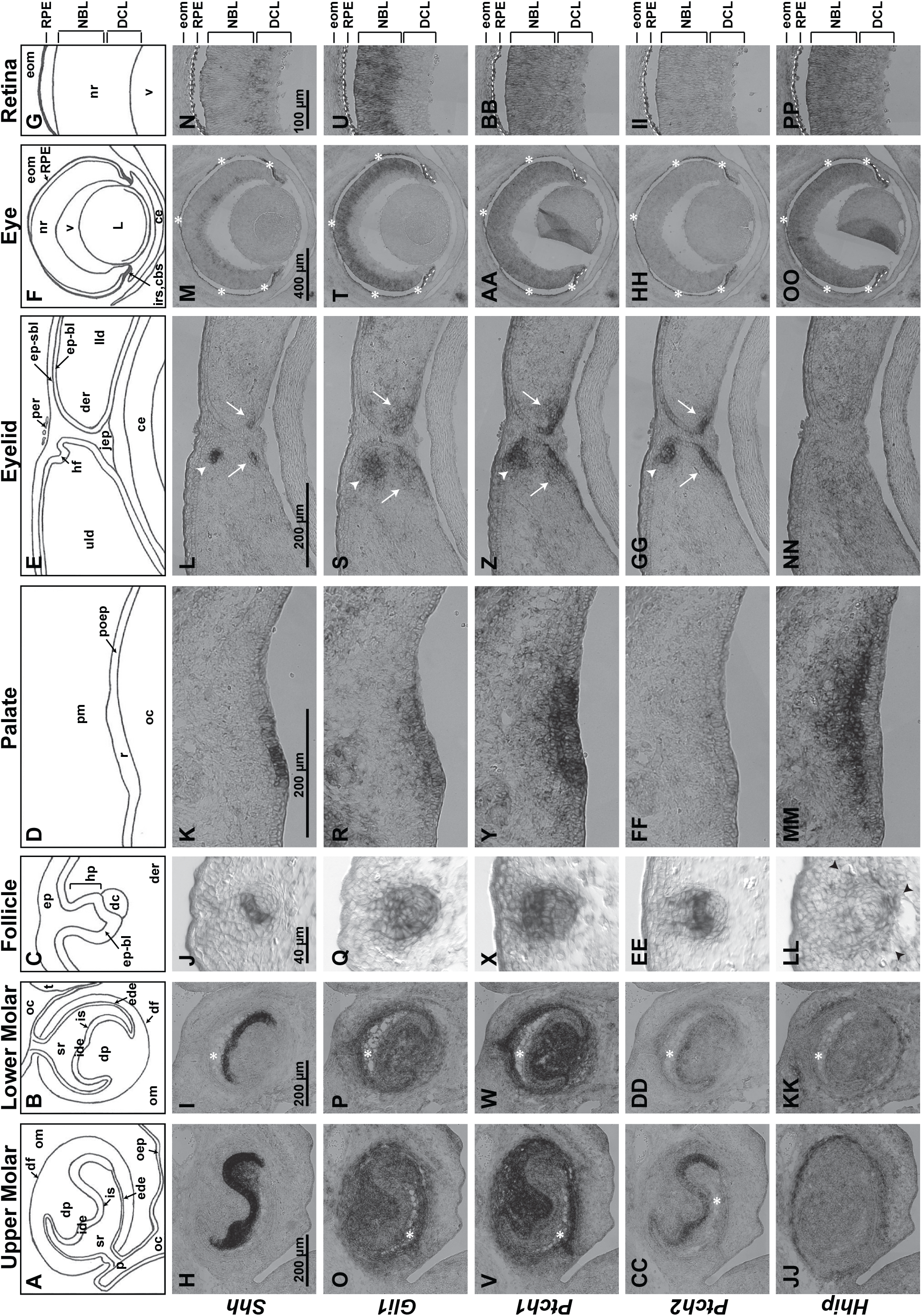
Close up comparisons of expression patterns. (A, B) Schematics of upper (A) and lower (B) molars with buccal side to the left, lingual side to the right. (C-G) Schematics of stage 3 hair follicle (C), palate (D) eyelid (E), eye (F), and retina (G). (H-PP) Expression patterns of *Shh* (H-N), *Gli1* (O-U), *Ptch1* (V-BB), *Ptch2* (CC-II), and *Hhip* (JJ-PP) in each structure. See abbreviations list for descriptions. White asterisks indicate histological artifacts where tissues are lacking. The asterisks for the eye structures also indicate the pigmentation of the RPE and is not mRNA staining. Black arrowheads in LL point to *Hhip* in the condensing mesenchyme surrounding the follicle. White arrows in eyelid panels point to the eyelid signaling field, and the arrowhead denotes a hair follicle. White dashed lines in eye panels denote the iris stroma (below line). White dashed lines in retina panels denote the RPE (below lines) and extraocular mesenchyme including the condensing scleral mesenchyme (above lines).

To better resolve the spatial characteristics of gene expression, Figure 2 shows the expression patterns at scales appropriate for each tissue. Illustrations for each structure are presented (Fig. 2A-G), with specific anatomical and gene expression descriptions provided in the supplemental text. As above, our focus here is to compare gene expression patterns across the structures.

In general, *Shh* expression was restricted to epithelial tissues within the molars, hair follicles, palatal rugae, and eyelids (Fig. 2H-L). The neural retina is primarily composed of cells from the neuroepithelium but *Shh* expression was similarly segregated, in this case, to the differentiated cell layer (DCL) where the RGCs are located (Fig. 2M,N). By and large, *Shh* expression was robust relative to the size of the tissue except in the retina, where expression was disproportionately lower.

*Gli1* and *Ptch1* exhibited largely overlapping patterns of expression (Fig. 2O-AA). Both were expressed throughout epithelial and mesenchymal tissues. Interestingly, mesenchymal tissues stained more strongly for *Gli1* and *Ptch1* in the molars and hair follicles (Fig. 2O-Q,V-X), while epithelial staining was stronger in the palate and eyelids (Fig. 2R,S,Y,Z). In the neural retina, *Gli1* expression overlapped with that of *Ptch1* in the neuroblast layer (NBL; Fig. 2T,U,AA,BB). *Gli1* was not detected in the DCL whereas *Ptch1* extended into the DCL.

*Ptch2* expression overlapped with *Shh* but was expressed more broadly (Fig. 2CC-EE,GG), consistent with earlier reports (20, 29). Two exceptions are the palatal rugae and retina where *Ptch2* expression were not detected (Fig. 2FF,HH,II). Since *Ptch2* is reliably detected in the retina by transcriptomic and RT-PCR based methods ((28), personal observation (XL and EML), the lack of detection here suggests low and potentially broad expression.

*Hhip* was expressed in a narrow band in the mesenchyme surrounding the molars at the outer edge of *Ptch1* and *Gli1* expression, at a distance from *Shh*-expressing cells (Fig. 2JJ,KK). Although not as distinct as in the molars, *Hhip* was expressed in the condensing mesenchyme surrounding the epithelial compartment of the hair follicle (Fig. 2LL; arrowheads in Fig. 1F denote additional follicles). *Hhip* expression within the palate exhibited a graded and robust pattern that was strongest in the palatal mesenchyme (pm) immediately adjacent to the *Shh*-expressing ruga (r). Expression in the eyelid was too low to assess (Fig. 2NN). As with *Ptch1, Hhip* expression in the retina extended across both the NBL and DCL in a graded manner that was strongest in the NBL.

## Discussion

Through direct comparative analysis, the expression patterns of several feedback regulators of Hh signaling were assessed. To first address the question that motivated this study, we found that the abundance of *Shh* mRNA is comparatively low in the retina, but pathway activity, as assessed by *Gli1* expression, is robust and on par with other tissues. This suggests tissue-specific differences in how robust signaling is achieved, and selective utilization of the negative feedback factors is one possibility. Supporting this, we observed nonoverlapping expression of *Ptch2* and *Hhip*, even in structures that express both. Thus, in addition to providing a mechanism to prevent overactive signaling, the utilization of specific feedback inhibitors could contribute to more efficient Hh signaling at lower levels of ligand expression and vice versa.

Of the three negative regulators, the expression pattern for *Ptch1* was most similar to *Gli1*. Although this makes it the least likely to have a tissue-selective role, it does make it the most reliable of the negative regulators to mark the field of active signaling. This is not surprising since Ptch1 is required in the majority of tissues for ligand-dependent signaling (14). Subtle differences, however, in its expression levels whether quantitative or spatial, or in the localization or modification of Ptch1 protein, could contribute to tissue-specific influences on signaling (30).

*Ptch2* and *Hhip*, however, exhibited unique expression characteristics. In the molars and hair follicles, their expression domains marked the two ends of the signaling field, with *Ptch2* closest to the source of ligand and Hhip expressed at the outermost extent of signaling. For the remaining structures, only *Ptch2* was expressed in the eyelid and only *Hhip* was expressed in the palate and retina, at least as could be assessed here. Although *Ptch2* and *Hhip* are both negative feedback regulators and act at the level of ligand availability, their differential utilization could account for differences in signaling efficiency across the structures. For example, if the retina is most efficient at Hh signaling as suggested, could *Hhip* have a role in this? How this might occur is not clear but there are differences in how Ptch2 and Hhip regulate ligand availability. Whereas both are found on the cell membrane where they bind and remove Shh ligand by endocytosis, Hhip also exists in a secreted form and sequesters ligand that remains extracellular. This could keep ligand intact, releasing it for signaling at a later time or in another location. Thus, Hhip could also have a supportive role in Hh signaling.

### Conclusions

This study reveals similarities and differences in the spatial expression patterns of *Shh, Gli1, Ptch1, Ptch2* and *Hhip* in 6 anatomical structures where Hh signaling occurs simultaneously. The expression patterns of *Gli1* and *Ptch1* suggest similar levels of signaling across structures with different levels of *Shh* expression. The expression patterns of *Ptch2* and *Hhip* suggest different roles in controlling the level of signaling in each tissue.

## LIMITATIONS

*in situ* hybridizations using colorimetric detection is qualitative in nature and does not allow for precise measurements of mRNA expression levels. Another limitation is the indirect nature of using gene expression as indicators of ligand availability or signaling activity. To test our hypothesis that differences in *Ptch2* and *Hhip* utilization contribute to qualitatively similar levels of *Gli1* expression and pathway activity in different anatomical structures would require functional perturbation experiments such as overexpression and knockdown of *Ptch2* and *Hhip* and evaluation of ligand availability for each structure.

## Abbreviations

## Molars

ide: internal dental epithelium;
ede: external dental epithelium;
is: intermediate stratum;
sr: stellate reticulum;
dp: dental papilla;
df: dental follicle;
p: pedicle;
oc: oral cavity;
om: oral mesenchyme;
oep: oral epithelium;
t: tongue.

## Hair follicles

ep: epidermis;
ep-bl: basal layer of epidermis;
hp: hair peg;
dc: dermal condensate;
der: dermis.

## Palatal rugae

r: palatal rugae;
poep: palatal oral epithelium;
pm: palatal mesenchyme;
oc: oral cavity.

## Eyelid

uld: upper lid;
lld: lower lid;
ep-sbl: suprabasal layer of epidermis;
ep-bl: basal layer of epidermis;
der: dermis;
jep: junctional epithelium;
per: residual periderm;
hf: hair follicle;
ce: corneal epithelium.

## Eye and retina

nr: neural retina;
RPE: retinal pigmented epithelium;
v: vitreous;
L: lens;
ce: corneal epithelium;
pom: periocular mesenchyme;
irs: iris stroma;
cbs: ciliary body stroma;
NBL: neuroblast layer;
DCL: differentiated cell layer;
eom.: extraocular mesenchyme.

## Acknowledgements

We thank Valerie Wallace for the plasmids used for RNA probe synthesis and Robert Marc and Bryan Jones for use of the Syncroscan-equipped Leica DMR upright microscope and imaging assistance.

## Authors’ contributions

EML and CLS conceived and wrote the study. CLS performed experiments. XL sequenced and mapped all probes. EML, CLS, and XL contributed to data analysis, figure preparation, and manuscript editing.

## Funding

This work was supported by funds from the NIH to EML (NEI R01-EY013760), a predoctoral traineeship to CS (NIGMS T32-GM007464), and with unrestricted funds to the John A. Moran Eye Center and the Vanderbilt Eye from Research to Prevent Blindness, Inc.

## Availability of data and materials

No additional datasets are associated with this study. Plasmids are available upon request to the corresponding author.

## DECLARATIONS

## Ethics approval and consent to participate

Consent not applicable. Animal use and care were conducted in accordance with protocols approved by the University of Utah Institutional Animal Care and Use Committee.

## Consent for publication

Not applicable.

## Competing interests

The authors have no competing interests to declare.

## FIGURE LEGENDS

**Supplemental Figure 1: In situ mRNA hybridization probe templates**. cDNA inserts were sequenced from each end, aligned with BlastN, and mapped onto their respective NCBI reference sequence using NCBI’s Sequence Viewer 3.24.0 (https://www.ncbi.nlm.nih.gov/tools/sviewer/)

**Supplemental Text: Anatomical descriptions of each structure and gene expression patterns**.

## Notes

### Competing Interest Statement

The authors have declared no competing interest.

