## Supplementary figures and images for "Expression of Sonic Hedgehog and Pathway Components in the Embryonic Mouse Head: Anatomical Relationships Between Regulators of Positive and Negative Feedback"

### Supplemental Figure 1

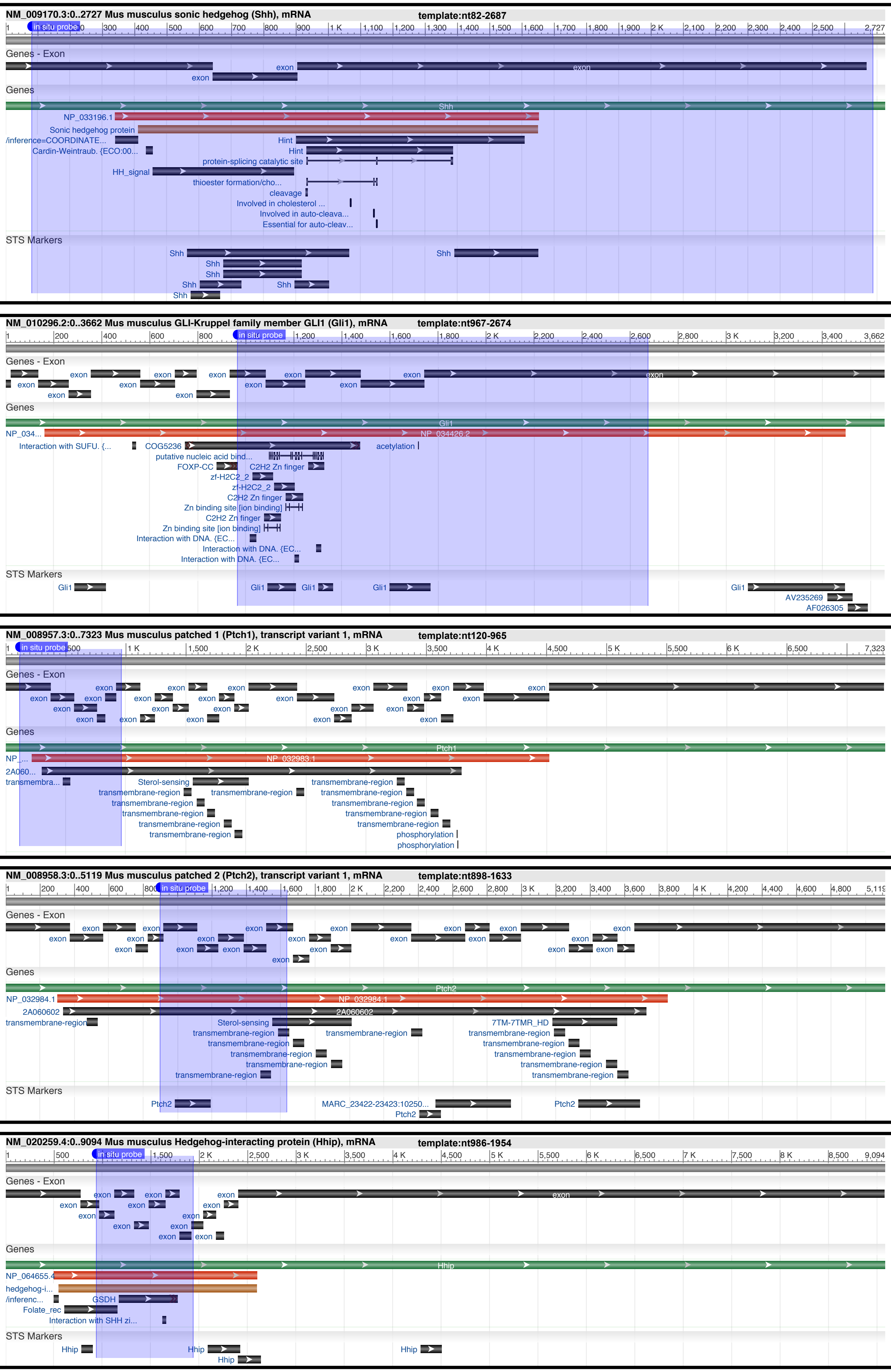
