## Supplemental Text for "Expression of Sonic Hedgehog and Pathway Components in the Embryonic Mouse Head: Anatomical Relationships Between Regulators of Positive and Negative Feedback"

This supplement provides detail on the anatomical organization of each tissue and how the expression of each gene is expressed within each anatomical framework. The abbreviations used in the illustrations in Figure 2 are provided here at first mention and in the abbreviations list in the main article.

### *Molars*

Odontogenesis of molars involves the budding of thickened oral epithelium (oep) and subsequent interactions with condensing neural-crest-derived mesenchymal cells (ectomesenchyme). By E15.5, the upper and lower molars have reached the late cap stage (Fig. 2A,B). The oral epithelium has undergone drastic morphological changes and histodifferentiation, giving rise to the internal and external dental epithelium (ide/ede, respectively), intermediate stratum (is), and stellate reticulum (sr), which connect to the oral epithelium via the pedicle (p). Condensation of the ectomesenchymal cells is the process of establishing the dental papilla (dp) and dental follicle (df).

Except where noted, our observations are consistent with prior characterizations (reviewed in (1)). *Shh* expression in the late cap stage is restricted to the epithelial components of both the upper and lower molars (Fig. 2B,H). Strong expression is observed within the internal dental epithelium and adjacent intermediate stratum. *Shh* expression also extends into varying portions of the stellate reticulum. In the upper molars, expression into the stellate reticulum is extensive, but in the lower molars is largely restricted to the intermediate stratum. Interestingly, expression in these epithelial tissues is more extensive on the lingual (tongue) side of both the upper and lower molars. Similar to *Shh*, *Ptch2* transcripts are detected throughout the internal and external dental epithelium (Fig. 2CC,DD). Strongest expression is observed within the internal dental epithelium and extends beyond the range of *Shh* expression, particularly on the buccal (cheek) side. In contrast, expression within the external dental epithelium is weak. Weak *Ptch2* signal is also observed within the intermediate stratum and stellate reticulum. This expression is not uniform and extends beyond the range of Shh expression. Contrary to previous reports (reviewed in Cobourne and Sharpe, 2005), *Ptch2* is also weakly detected within the condensing ectomesenchyme of the forming dental papilla and follicle. In the upper and lower molars, *Ptch1* is expressed throughout both epithelial and mesenchymal components (Fig. 2V,W). Strongest expression is observed in the condensing ectomesenchyme, including that forming the dental papilla and follicle, and in the most peripheral stellate reticulum and pedicle. Robust expression is also found in the internal and external dental epithelium, intermediate stratum, and distal stellate reticulum. Strong expression of *Hhip* transcripts is restricted to a narrow band of peripheral mesenchyme, including the dental follicle, which encompasses the developing molars, many cell diameters from *Shh*-expressing cells (Fig. 2JJ,KK). Faint *Hhip* expression is also detected throughout the remaining mesenchymal and epithelial components of the developing upper and lower molars. *Gli1* expression (Fig. 2O,P) mirrors that of *Ptch1*. Interestingly, *Gli1* expression fails to extend into the oral mesenchyme (om) much beyond the extent of strongest *Hhip* expression in the forming dental follicle. Because *Gli1* provides a reliable readout of active Hh signaling, expression of *Gli1* transcripts reveals Hh pathway activation throughout the epithelial and mesenchymal tissues of the developing molars.

***Hair follicles***

Like molars, hair follicles (HF) arise through a series of interactions between the epidermis (ep) and underlying mesenchyme. The basal layer of the epidermis thickens and grows downward toward condensing dermal mesenchyme. By E15.5, most tylotrich pelage hair follicles (2, 3) have reached stage 3 of hair follicle morphogenesis (based on morphological classification described in (3-5)). At this stage, the follicle, now termed a hair peg (hp), consists of a solid column of epidermally-derived epithelial cells with a concave end that partially or wholly encompasses the mesenchymal dermal condensate (dc), which will form the future dermal papilla (Fig. 2C).

Expression of *Shh* within the stage 3 hair follicle is restricted to the distal end of the epithelial downgrowth, within the concave basal border cells that contact the dermal condensate (Fig. 2J). *Ptch2* is similarly expressed in the distal epithelium of the hair peg (Fig. 2DD), overlapping the region of *Shh* expression. However, weaker *Ptch2* expression extends beyond the *Shh*-expressing region, up the outer walls of the hair peg. *Ptch1* is expressed in both the epithelial and mesenchymal components of the developing hair follicle (Fig. 2X). Strongest expression is observed in the distal half of the epithelial-derived hair peg, both in the basal layer epidermal cells, which comprise the outer layer cells of the hair peg, and cells of the interior, as well as uniformly through the dermal condensate. The most proximal aspect of the hair peg only weakly expresses *Ptch1*. Weak *Ptch1* expression also extends slightly into the surrounding noncondensed dermal mesenchyme directly adjacent to the hair peg and dermal condensate. Expression of *Hhip* is strongest in a narrow region of the noncondensing dermal mesenchyme surrounding the dermal condensate and hair peg (Fig. 2LL). Weak expression is also observed in the peripheral cells of the dermal condensate and inner portions of the hair peg. *Gli1* expression (Fig. 2Q) again mimics that of *Ptch1* and is largely restricted within the border of high *Hhip* expression. *Gli1* expression reveals active Hh reception and signaling in both the mesenchymal and epithelial components.

Expression of *Shh* and *Ptch2* at this stage (stage 3) is consistent with their expression patterns reported for other stages (stages 0-2 and stages 4-5; (6). The expression patterns of *Ptch1* and *Gli1* at this stage are also consistent with reports for earlier stages (0-2), but are much broader than that for Stages 4-5 (6).

***Palatal rugae***

Palate development is a complex process involving elevation, midline contact and eventual fusing of the palatal shelves above the tongue (7). In the murine palate at E15.5, fusion of the palatal shelves is mostly complete, and the medial edge epithelial seam is undetectable (Fig. 2D).

At this rather late stage of palate development, expression of Hh pathway components in the palatal tissues is largely restricted to the developing rugae (7). However, some pathway components are also expressed in the developing palatine bones in response to known Ihh signals in this structure (7-9).

Expression of *Shh* transcripts in the palate is restricted to the small, thickened regions of palatal oral epithelium (poep) known as rugae (r) (Fig. 2K). Unlike the other epithelial/mesenchymal organs described here, a clear *Ptch2* signal is undetectable in the palatal oral epithelium (Fig. 2FF), consistent with previous reports (7). *Ptch2* expression was also not observed in the palatal mesenchyme (pm). In previous reports, *Ptch2* was weakly detected only in anterior palatal mesenchyme (7). *Ptch1* is strongly expressed by the thickened palatal oral epithelium in a region that overlaps with but extends beyond that of *Shh* expression (Fig. 2Y). Robust *Ptch1* expression is also observed extending into the palatal mesenchyme immediately adjacent to the palatal oral epithelium in a gradient fashion. *Hhip* is strongly expressed in the palatal mesenchyme adjacent to the palatal oral epithelium of the developing rugae (Fig. 2MM), in a large region that overlaps and extends beyond that of *Gli1* and *Ptch1* expression. This mesenchymal expression is also graded in nature, with the strongest expression closest to the oral epithelium. *Gli1* transcripts are again expressed in a pattern similar to that observed for *Ptch1* in rugae of the palatal oral epithelium and adjacent mesenchyme, although slightly more restricted in range and less uniform within the mesenchymal component (Fig. 2R). Thus, epithelial-derived Shh activates Hh signaling in both the palatal epithelial and adjacent mesenchymal compartments of the palate.

***Eyelid***

During eyelid development, the lid primordia emerge as folds from the epidermis and dermis (der) surrounding the eye, which then grow to extend over the corneal epithelium (ce) of the eye until they meet and fuse, only to reopen later. By E15.5 in the mouse, the upper and lower eyelids (uld/lld, respectively) have extended over the corneal epithelium and fused at approximately the center of the eye (Fig. 2E).

Expression patterns of *Shh*, *Ptch1* and *Ptch2* have been reported in the eyelid prior to fusion, during the extension phase (10), and appear to be restricted to the basal layer of the eyelid epithelium (ep-bl). Following fusion, expression of *Shh*, *Ptch1* and *Ptch2* continues in the basal layer of the eyelid epithelium originating from both the upper and lower lid primordium (Fig. 2L,Z,GG). *Shh* expression is restricted to a small patch of basal layer eyelid epithelium on the corneal side (Fig. 2L). *Ptch2* expression is similarly restricted to the basal layer eyelid epithelium but exhibits a broader expression territory that overlaps and extends beyond the region of *Shh* expression (Fig. 2GG). *Ptch1* is also strongly expressed within a similar range of basal layer eyelid epithelium (Fig. 2Z). However, unlike the epithelium-restricted expression observed in the extension stage (10), *Ptch1* expression clearly extends into the adjacent mesenchyme of the eyelid tip, although at weaker levels. *Hhip* is only faintly detected in the mesenchyme of the eyelid tip (Fig. 2NN). *Gli1* is expressed in both the basal layer eyelid epithelium and adjacent mesenchyme of the eyelid tip (Fig. 2S) in a range comparable to that of *Ptch1* and reveals active Hh signaling in these tissues.

***Ocular tissues***

Eye development involves a series of complicated morphogenetic processes in which the neural retina (nr), retinal pigmented epithelium (RPE), and optic stalk derive from the neuroectoderm and come to surround the surface ectoderm-derived lens capsule in a cup-like fashion. By E15.5, the lens vesicle (L) has detached from the overlying surface ectoderm and the remaining surface ectodermal cells together with migrating mesenchymal cells have condensed to form the layers of the corneal epithelium. In the optic cup, the RPE monolayer surrounding the neural retina has become pigmented (Fig. 2F). Retinal neurogenesis is underway, and the neural retina contains two morphologically distinct layers: a basal differentiated cell layer (DCL) containing nascent retinal ganglion cells and amacrine cells and an apical neuroblast layer (NBL) primarily composed of progenitors but interspersed with nascent horizontal cells and cone photoreceptors (Fig. 2G).

Expression of Hh pathway components in the eye at E15.5 is restricted to the developing retina, cells of the developing choroid and sclera surrounding the RPE and differentiating stromal cells of the iris (irs) and ciliary body (cbs). In the neural retina, *Shh* expression is restricted to the differentiated cell layer (Fig. 2M,N), consistent with its production by retinal ganglion cells (11). Similar to the palate, *Ptch2* staining is too close to background levels to make a definitive assessment (Fig. 2HH,II). *Ptch2* is readily detected by RT-PCR and RNA-sequencing at E15.5 (XL, EML, personal observations), so the difficulty to ascertain its pattern could suggest low but broad expression. Expression of *Ptch1* extends throughout both retinal layers, with slightly stronger expression observed in the neuroblast layer (Fig. 2AA,BB). *Hhip* is detected throughout the neural retina, with slightly stronger expression observed in the neuroblast layer (Fig. 2OO,PP). In contrast, expression of *Gli1* is definitively restricted to the neuroblast layer (Fig. 2T,U), consistent with the established roles of Shh and Hh signaling in the proliferation and cell fate decisions of retinal progenitor cells (11).

A narrow band of *Gli1, Ptch1* and *Hhip* expression is also observed outside the retina in the scleral condensation of the extraocular mesenchyme (eom) observed above the dotted lines in Fig. 2U, BB, and PP, which mark the boundary with the RPE (below line). These are in response to Indian Hedgehog (Ihh) produced by endothelial cells of the developing choroidal vasculature, a layer of cells situated between the RPE and scleral condensation immediately above the dotted line (12-14). Analyses of Hh target gene expression in ocular tissues of *Ihh*-null mice and conditional *Shh* mutants reveal that the range of neuron-derived Shh action is restricted to the neural retina, while the range of Ihh action is restricted to the RPE and periocular mesenchyme (12, 14). Although difficult to observe at the scale presented, *Gli1* and weak *Ptch1* expression are also detected in the stroma of the iris, located below the dotted lines in Fig. 2T and AA. The lack of detectable *Shh* expression in this region (Fig. 2M) suggests this expression may also be in response to *Ihh*. Supporting this, *Ihh* was detected in the adult iris of mouse and newt, together with *Ptch1* and *Ptch2* (15), and *Gli1* expression was not reported in the iris stroma in *Ihh* mutant mice (12, 14).
